# DEVELOPMENT OF PROTOCOLS FOR THE CULTIVATION OF WORMWOOD (*ARTEMISIA SALSOLOIDES*)

**DOI:** 10.1101/2025.03.31.646446

**Authors:** Vasiliy A. Chokheli, Anatoly S. Azarov, Victoria S. Petrenko, Mark O. Belyaev, Klimentiy A. Tulupov, Christina A. Tsymerskaia, Svetlana N. Sushkova

## Abstract

As part of the work, the key factors influencing the growth and development of *Artemisia salsoloides* culture have been studied, and nutrient media have been developed and optimized for plant conservation. The optimal medium for the cultivation of *A. salsoloides* is MS + 2 mg/L MT. Viable suspensions were obtained. The MS nutrient medium and the BM1k nutrient medium developed by us are suitable for wormwood callusogenesis. The growth curves of cell suspensions have a standard S-shaped curve. In the *A. salsoloides* plants we studied, key genes for the synthesis of artemisinin-like compounds (ADS, CYP71AV1, DBR2) were found, except for the CPR gene.

## INTRODUCTION

Along with the widespread use of *ex situ* conservation methods for threatened plant species, the International Botanical Gardens Plant Protection Program (2000) provides for the creation of *in vitro* collections of plant material. Thanks to such collections, it is possible to cultivate plant species whose *ex situ* cultivation is difficult or impossible (Pence, 1999; Andreev, Gorbunov, 2000). The attractiveness of creating such collections lies in the absence of the need to remove large amounts of plant material from natural habitats (Engelmann, 2011; Laslo et al., 2011).

Despite the widespread use of the micropropagation method and its long-standing use in biotechnology (Butenko, 1990), the development of microcloning techniques for rare and endemic plant species is an incentive for the emergence of new techniques in this field (Chokheli et al., 2020). Thus, today, the peculiarities of the chemical composition of the soils of plant communities where a rare plant species grow, which is necessary for introduction into culture, are increasingly being considered (Fay, 1992).

Micropropagation of many valuable endangered plants is performed in bioreactors (Ahmadian et al., 2017). Studies using bioreactor systems have been performed for many plant families with many rare species: *Orchidaceae* Juss. (Yoon et al., 2007), *Plantaginaceae* Juss. (Ahmadian et al., 2017), *Araceae* Juss. (Arano-Avalosa et al., 2020).

*Artemisia* L. (family *Compositaceae*, formerly called *Asteraceae*) is the largest genus of the tribe *Anthemideae*, comprising more than 500 species distributed almost worldwide in both temperate and tropical regions. Extracts from some species of the genus *Artemisia* are actively used in medicine as antiparasitic agents. An extract from *Artemisia annua* is used to produce artemisinin, an active substance that suppresses the development of malarial plasmodium (Muangphrom et al., 2016).

One of the promising methods for the conservation of rare and endangered *Artemisia* species is the creation of suspension cultures that enable long-term and effective conservation of the gene pool and the production of biologically active substances with many valuable pharmacological properties. However, to achieve optimal results, it is necessary to develop methods for optimizing nutrient media that ensure the highest productivity and quality of suspension culture.

The purpose of this study is to develop protocols for cultivating *Artemisia salsoloides* plants.

## MATERIALS AND METHODS

### Plant material

The objects of our research are plants of the wormwood (*A. salsoloides*). Wormwood seeds were collected from the collection “Rare and Endangered plants of the Rostov Region” of the Botanical Garden of the SFedU in 2023. The laboratory experiment was conducted in 2024.

*A*.*salsoloides* belongs to the angiosperm division (*Magnoliophyta* (*Angiospermae*), dicotyledonous class (*Magnoliopsida*), Astrocolor order (*Asterales*), and *Asteraceae* family. *A*.*salsoloides* is included in the “Red List of the Russian Federation”, “Red List of the Krasnodar Territory”, “Red List of the Rostov region”. The species is part of the sublittoral communities of the seashore. It occurs on sandy-shell and limestone substrates (cl. *Ammophlietea, Onosmo polyphyllae-Ptilostemonetea*). Xerophyte. Obligate calcephylus, growing on outcrops and rocky steppes with underlying carbonate rocks (Red List of the Russian Federation, 2008; Red List of the Russian Federation, 2014).

### Sterilization of planting material

The seed material was sterilized according to a proven method (Chokheli et al., 2020) by washing the seeds in a mixture of 70% ethyl alcohol solution and 3% hydrogen peroxide solution in a 1:1 ratio for 10 minutes, followed by washing the diaspores in distilled water for 10 minutes 4 times.

### Introduction to in vitro culture

The seeds were germinated on Murashige-Skoog nutrient medium (Murashige, Skoog, 1962) with the addition of growth regulators methatopolyn (MT), 6-benzylaminopurine (BAP) and forchlorfenuron (N- (2-chloro-4-pyridyl) -N’-phenylurea) (CPPU) (Table 1).

**Table 1.** Variants of nutrient media for cultivating *Artemisia salsoloides* plants *in vitro*.

|  | **BAP, mg/L** | **MT, mg/L** | **CPPU, mg/L** |
| --- | --- | --- | --- |
| **MS** | 0.2 | 0.5 | 0.1 |
|  | 0.5 | 1 | 0.2 |
|  | 0.8 | 1.5 | 0.3 |
|  | 1.2 | 2 | 0.4 |
|  | 1.5 | 2.5 | 0.5 |

The explants and plant passages were sterilized in a laminar flow box using instruments sterilized in a drying cabinet (scissors, tweezers, dissecting needles). For nutrient media, the pH is adjusted to 6.0 using a 1 M solution of potassium hydroxide KOH. The culture media was sterilized in an autoclave MLS-3751L (Sanyo) at a temperature of 121° C and a pressure of 1.5 atmospheres for 30 minutes. The plants were cultivated at a constant temperature of 25°C and a 16-hour photoperiod, with a light intensity of 3500 lux.

### Callusogenesis

The object of the study was the callus cultures of *A. salsoloides*. The cultures were grown on agarized MS nutrient medium and experimental BM1k medium by the surface method, at a temperature of 26 ± 1° C, in the dark for 5 days and then in the light at an illumination of 3800 lux and a temperature of 26 ± 1° C for up to 30 days. The compositions of MS and BM1k nutrient media are shown in Table 2. Explants to produce wormwood callus of *A. salsoloides* were parts of stems with leaves ∼ 5-8 mm in size, obtained from plants grown *in vitro* on MS medium.

**Table 2.** Ionic composition of media (mg/L)

| Ions | BM1k | MS |
| --- | --- | --- |
| K<sup>+</sup> | 509.72 | 781.66 |
| NH<sub>4</sub><sup>+</sup> | 135.16 | 371.68 |
| Ca<sup>2+</sup> | 211.73 | 119.72 |
| Mg<sup>2+</sup> | 53.24 | 36.48 |
| Na<sup>+</sup> | 6.00 | 4.61 |
| NO<sub>3</sub><sup>-</sup> | 1734.38 | 2443.29 |
| SO<sub>4</sub><sup>-</sup> | 222.98 | 153.83 |
| Cl<sup>-</sup> | 47.62 | 212.50 |
| PO<sub>4</sub><sup>3-</sup> | 174.47 | 118.64 |

The media were prepared in a standard way and sterilized in an autoclave at a temperature of 121°C for 20 minutes.

Explants were transferred into test tubes, one explant at a time. And they were placed in a thermostat at a temperature of +26°C. Callus growth induction was performed in the dark.

Two days later, after replanting the culture on a fresh nutrient medium, the first sample was taken to determine the raw mass of the callus. The callus culture was separated from the agarized nutrient medium and weighed on an analytical scale. Next, the raw biomass was dried at a temperature of 105°C and then weighed again to determine the dry weight.

On days 6, 9, 12, 16, 20, 25 and 30, identical procedures were performed to determine the wet and dry weight of callus cultures.

### Maintenance of cell suspension

To culture the suspension of *A. salsoloides*, liquid nutrient media were used: MS and 1/2 MS in the NAA 2 mg/L + BAP 0.2 mg/L and 2.4-D 2 mg/L + BAP 0.2 mg/L variations. The cultivation of suspensions took place at a 16/8-hour photoperiod.

### Genetic analysis

Total DNA was isolated from homogenized samples in 70 mg batches, which were previously disinfected and treated with a weak NaClO solution. DNA isolation was performed using a commercial kit (Syntol, Russia) “Sorb-GMO-B” (CTAB + Sorbent).

The DNA concentration was determined by standard methods in accordance with the manufacturer’s protocol on an SP-UMUV100 spectrophotometer (Scitek, China).

Specific primers for the main artemisinin synthesis genes, *ADS, CYP71AV1, CPR*, and *DBR2*, were used for genetic analysis. The primers were selected based on literature data (Firsov et al., 2021). The sequence of primers is shown in Table 3. The melting point (Tm) of the primers was determined by the formula:

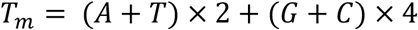

When setting the temperature gradient, we tried to select temperatures such that Tm was in the middle of the gradient.

**Table 3.**
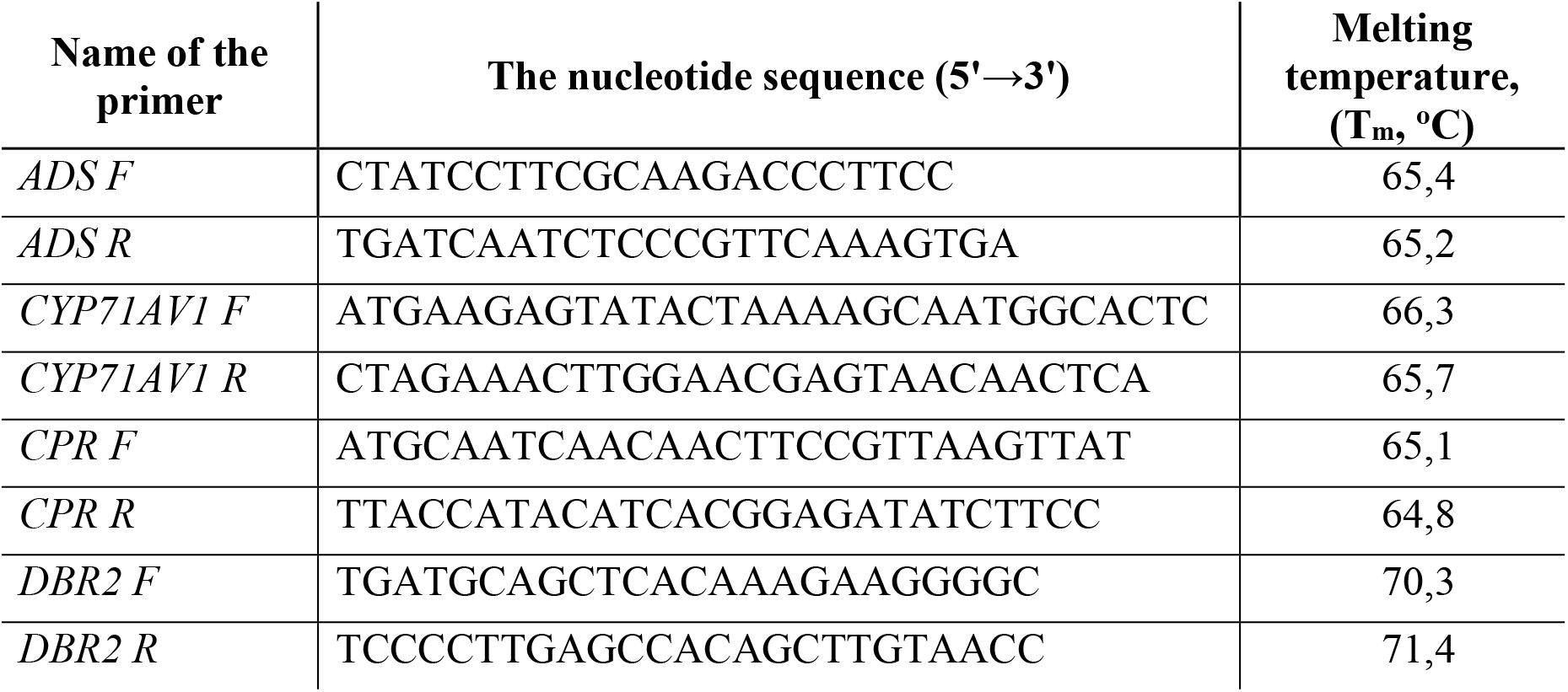
Characteristics of primers.

The PCR mixture per sample consisted of: H_2_O (DD) – 12.6 μL; 2.5mM dNTP – 2.5 μL; 10xTaq Turbo Bufer – 2.5 μL; MgCl_2_ (25 mM) – 3.0 μL; HS Taq polymerase (5 U/μL) – 0.4 μL; forward primer – 1 μL and a reverse primer (concentration of 10 pM/μL) − 1 μL; a DNA sample - 2 μL. The total volume of the mixture was 25 μL. Amplification was performed in a T100 Thermal Cycler (Bio-Rad, USA) and a C 1000 Thermal Cycler (Bio-Rad, USA). Amplification protocol: 1. 95 °C – 5:00 min.; 2. 94 °C – 0:30 s; 3. Ta °C – 0:30 s; 4. 72 °C – 2:00 min.; 5. 35 cycles points 2-4; 6. 72 °C – 5:00 min.; 7. Storage at 12 °C. The amplification was carried out with an annealing temperature gradient (Ta) of 54-65 °C.

The fragments were separated by electrophoresis in 1.5% agarose gel using 1×TBE buffer (Tris, Boric acid, EDTA) at a voltage of 80 V, 3 hours, power source − PowerPacBasic (BIO-RAD, USA). Tubes with amplicons were pre-mixed with 2 microns of the loading dye bromophenol blue + glycerin.

DNA staining was performed with SYBR Green I x78 dye (Lumiprobe, USA) at a ratio of 2 μL of dye per 5 μL of amplicons, and filming was performed in the GelDoc XR+ geldocumentation system (BIO−RAD, USA) with ImageLab software version 6.0 (BIO-RAD, USA). DNA fragment length markers 50+ bp, 100+ bp, and 1 kb+ bp DNA Ladder (Eurogen, Russia) were added 7 μL per well.

### Statistical processing

The quality of the nutrient medium can be assessed by the curves of cell growth and the number of cells in the volume of liquid. The measurements were carried out over 14 days. Every day, the number of cells per volume of liquid in the Goryaev chamber was evaluated in accordance with the method of Peskova et al. (2020). The number of cells in 1 mL of the suspension studied is calculated by the formula:

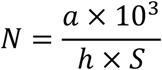

where **N** – is the number of cells in 1 mL of suspension; a is the average number of cells in the grid square; **10**^**3**^ – is the conversion coefficient [cm^3^] to [mm^3^]; **h** – is the depth of the chamber, mm; **S** – is the area of the grid square, mm^2^.

Growth indicators, including an increase in biomass, specific growth rate, growth index, the time of doubling of biomass, as well as the water content in cells were determined by calculation methods using the formulas given below (Kuznetsova et al., 2015). The formula for determining the increase in biomass (g) has the form:

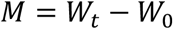

where **W**_**t**_ – is the final weight of biomass (g); **W**_**0**_ – is the initial weight of biomass (g). The formula for determining the specific growth rate (day–1) has the form:

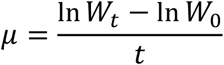

where **W**_**t**_ – is the final weight of biomass (g); **W**_**0**_ – is the initial weight of biomass (g), **t** – is the cultivation time (day). The formula for determining the growth index (relative units) has the form:

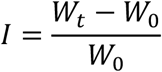

where **W**_**t**_ – is the final weight of biomass (g); **W**_**0**_ – is the initial weight of biomass (g). The formula for determining the doubling time of biomass has the form:

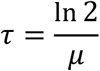

where **μ** – is the specific growth rate (day-1).

The formula for determining the water content in cells (%) has the form:

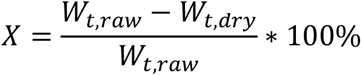

where **W**_**t**,**raw**_ – is the final weight of the raw biomass, **W**_**t**,**dry**_ is the final weight of the dry biomass.

Cytological preparations were prepared by the pressure method using the propionic-lacmoid method of staining the nuclei.

The significance of the differences was calculated according to the t-test (Lakin, 1990).

## RESULTS AND DISCUSSION

### Introduction to in vitro culture

The seeds were germinated on MS nutrient medium with the addition of BAP at a concentration of 0.5 mg/L. To accumulate the mass of *Artemisia salsoloides* explants, MS nutrient medium was used, the main task was to determine the growth regulator, which allows for the shortest possible time to increase plant mass. The data obtained are presented in Table 4.

**Table 4.**
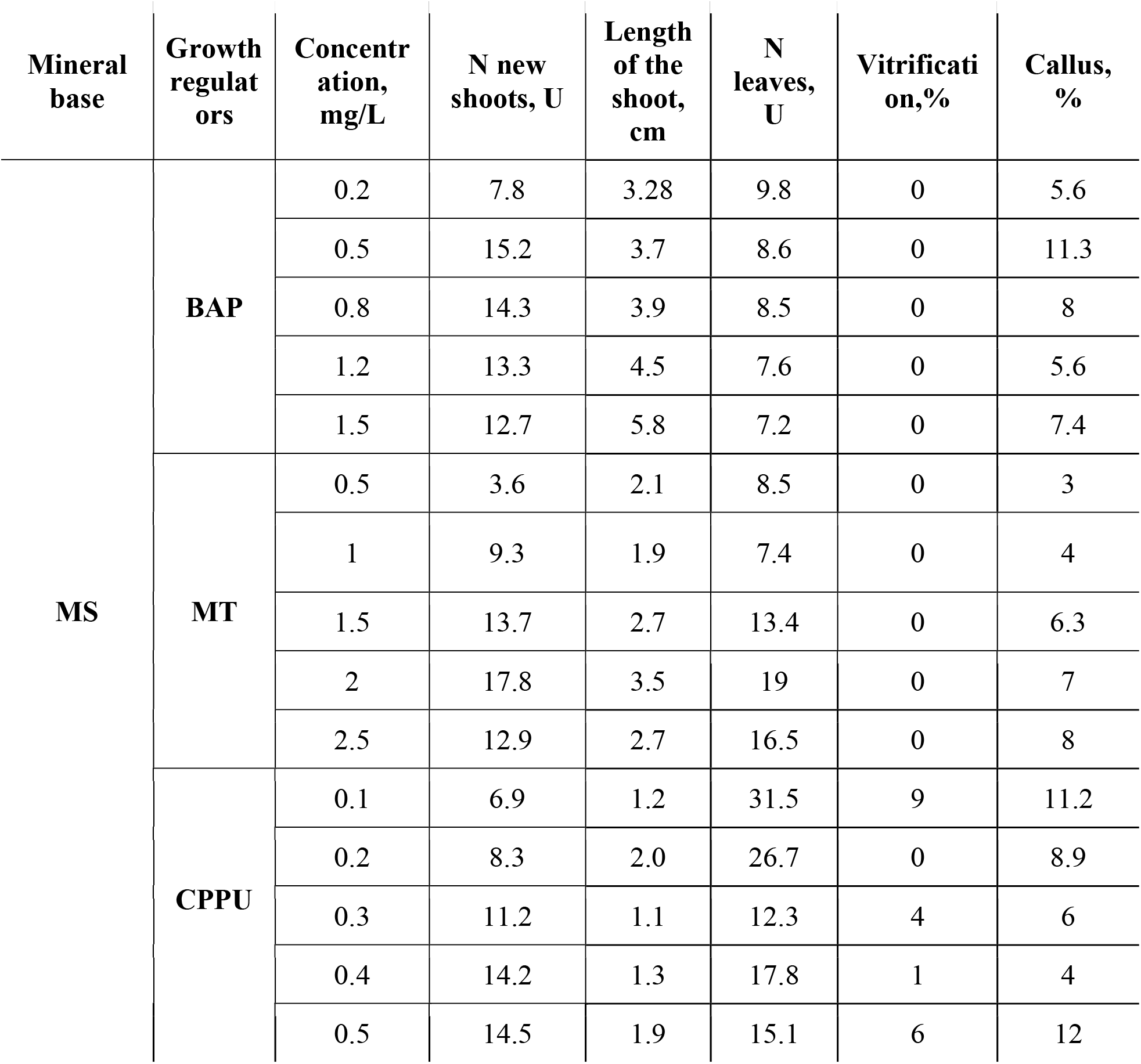
Growth characteristics of *A. salsoloides*.

As a result of the experiment, it was found that for the cultivation of *A. salsoloides*, the optimal medium is MS with the addition of MT at a concentration of 2 mg/L (Figure 1). This allowed us to achieve the highest multiplication coefficient (17.8 new shoots) and minimum vitrification (0%) and callus formation (7%).

**Figure 1.**
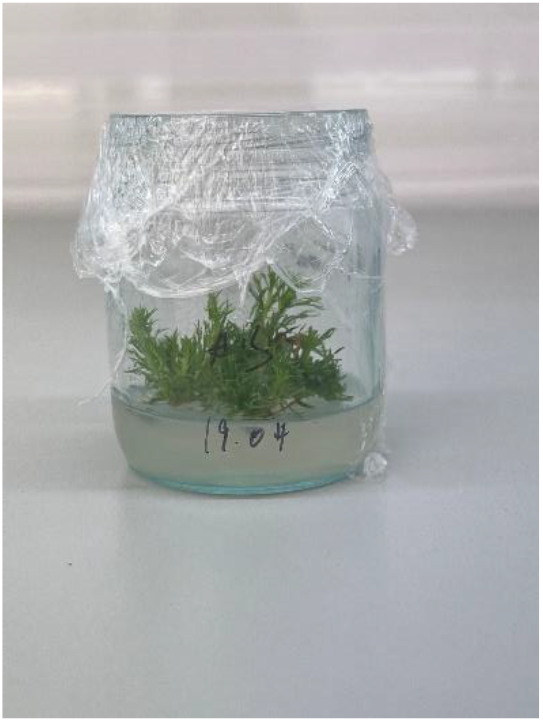
*A. salsoloides* on MS nutrient medium with addition of MT at a concentration of 2 mg/L.

The lowest multiplier coefficient was found for MS medium with the addition of CPPU at a concentration of 0.1 mg/L (6.9 new shoots).

We also compared the different concentrations of growth regulators used.

6-Benzylaminopurine (BAP) is a synthetic analog of the plant hormone cytokinin, which is widely used to stimulate the growth and development of plant cells. Research shows that BAP can positively influence the growth and development of plant cells, increasing their division and differentiation.

Various studies show that BAP affects the growth and development of cells at the root of the *Arabidopsis thaliana* plant. The researchers found that BAP stimulates the growth of root hairs and increases the number of cells in the root. The effect of BAP on the growth and development of leaf cells of the Solanum lycopersicum plant has also been confirmed. Researchers have found that BAP stimulates the growth of leaf cells and increases their size (Kumlay et al., 2015).

The optimal concentration for the cultivation of *A. salsoloides* using BAP is 0.5 mg/L. At the same time, the multiplication coefficient increases to this concentration. And with increasing concentration, the multiplier decreases, from 14.3 at a concentration of 0.8 mg/L to 12.7 at a concentration of 1.5 mg/L.

It is known that BAP in plant tissues is metabolized to stable toxic compounds – 6-benzylaminopurine-9-glycosides, which accumulate in tissues at the base of plants, inhibiting their further development (Bairu et al., 2007). MT is a less toxic phytohormone that improves further processes of root formation *in vitro* and acclimatization. MT enhances the processes of multiplication and rhizogenesis, does not cause tissue hydration, and increases the success of plant acclimatization (Werbrouck et al., 1995; Strnad et al., 1997).

The optimal concentration for the cultivation of *A. salsoloides* using MT is 2 mg/L. At the same time, up to this concentration, the multiplier coefficient increases from 3.6 at a concentration of 0.5 mg/L to 13.6 at a concentration of 1.5 mg/L. And with increasing concentration, the multiplier decreases.

Like TDZ, 4-CPPU is also a phenylurea with cytokinin activity and stimulating shoot growth and organogenesis in vitro, mainly due to its relative tolerance to endogenous cytokinine oxidases, a key enzyme in cytokinin degradation, as well as its ability to produce endogenous cytokinins (Arinaitwe et al., 2000; Faisal et al., 2019).

The optimal concentration for cultivating *A. salsoloides* using CPU is 0.5 mg/L. The multiplication coefficient is 14.5. At a concentration of 0.1 mg/L, the minimum multiplication coefficient is 6.9.

Thus, the optimal medium for the cultivation of *A. salsoloides* is MS + 2 mg/L MT.

### Callusogenesis

It is known that callus cells undergo a few developmental phases during growth: lag phase, exponential, stationary, and degradation phase (Kuznetsova et al., 2015). As a result of the structural heterogeneity of the callus tissue, a physiological imbalance of the cell population is also observed. Using such indicators as the growth index, specific rate and time of biomass doubling, it is possible to characterize the growth of callus crops. The data obtained during the study are shown in Tables 5 and 6.

**Table 5.**
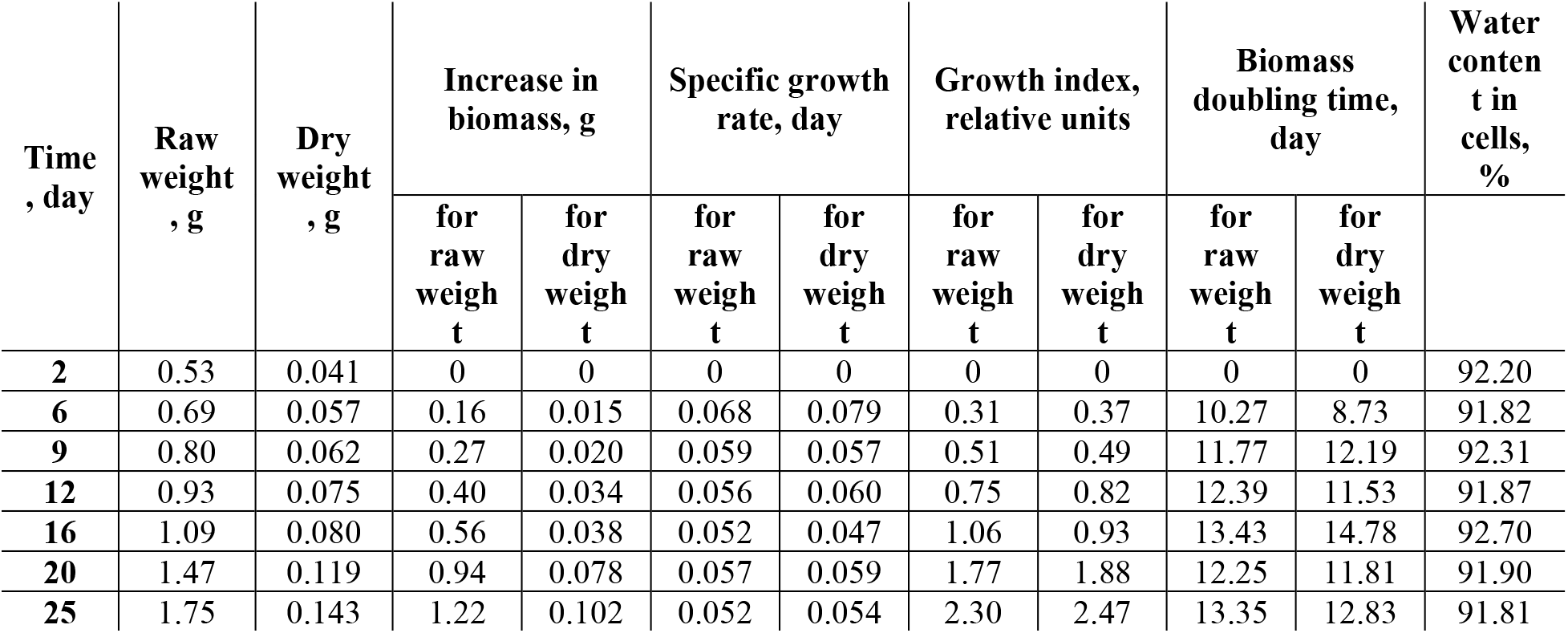
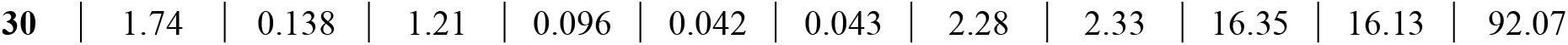
Dynamic growth rates of *A. salsoloides* calluses on MS medium (the arithmetic mean values of the results of three determinations with a deviation of no more than 5% by wet weight are given).

**Table 6.** Dynamic growth rates of *A. salsoloides* calluses on BM1k medium (the arithmetic mean values of the results of three determinations with a deviation of no more than 5% by wet weight are given)

| Time, day | Raw weight, g | Dry weight, g | Increase in biomass, g |  | Specific growth rate, day |  | Growth index, relative units |  | Biomass doubling time, day |  | Water content in cells, % |
| --- | --- | --- | --- | --- | --- | --- | --- | --- | --- | --- | --- |
|  |  |  | for raw weight | for dry weight | for raw weight | for dry weight | for raw weight | for dry weight | for raw weight | for dry weight |  |
| 2 | 0.48 | 0.044 | 0 | 0 | 0 | 0 | 0 | 0 | 0 | 0 | 90.90 |
| 6 | 0.65 | 0.059 | 0.17 | 0.015 | 0.077 | 0.073 | 0.36 | 0.34 | 9.02 | 9.46 | 91.03 |
| 9 | 0.77 | 0.073 | 0.29 | 0.029 | 0.067 | 0.073 | 0.60 | 0.67 | 10.32 | 9.46 | 90.50 |
| 12 | 0.87 | 0.087 | 0.39 | 0.044 | 0.060 | 0.069 | 0.82 | 1.00 | 11.57 | 10.01 | 90.01 |
| 16 | 1.01 | 0.088 | 0.53 | 0.044 | 0.053 | 0.050 | 1.10 | 1.01 | 13.08 | 13.92 | 91.30 |
| 20 | 1.26 | 0.107 | 0.78 | 0.063 | 0.054 | 0.050 | 1.62 | 1.45 | 12.95 | 13.94 | 91.50 |
| 25 | 1.50 | 0.139 | 1.02 | 0.095 | 0.049 | 0.050 | 2.12 | 2.17 | 14.01 | 13.81 | 90.75 |
| 30 | 1.47 | 0.126 | 0.99 | 0.083 | 0.40 | 0.038 | 2.07 | 1.89 | 17.30 | 18.26 | 91.42 |

The growth dynamics of callus crops have an S-shaped appearance, which is typical for most plant crops. The callus growth charts clearly show the intervals: the initial period of slow growth (lag phase) is from days 2 to 9, and the exponential growth phase is from days 9 to 20. The phase of growth deceleration from 20 to 25 days and the stationary phase from 25 to 30 days are obviously determined in *A. salsoloides* calluses on both media. For most crops, the growth cycle is 30 days, with the stationary phase occurring on an average of 20 days. According to the data on the accumulation of dry biomass, a general growth trend should be noted, however, due to the specifics of environments and stages of physiological development of tissues, differences in water availability and a non-linear dependence of the increase in dry biomass are observed over time. On the MS medium, the calluses are looser, more watered, and their production is promising for use in suspension culture to obtain secondary metabolites (Figure 2).

**Figure 2.**
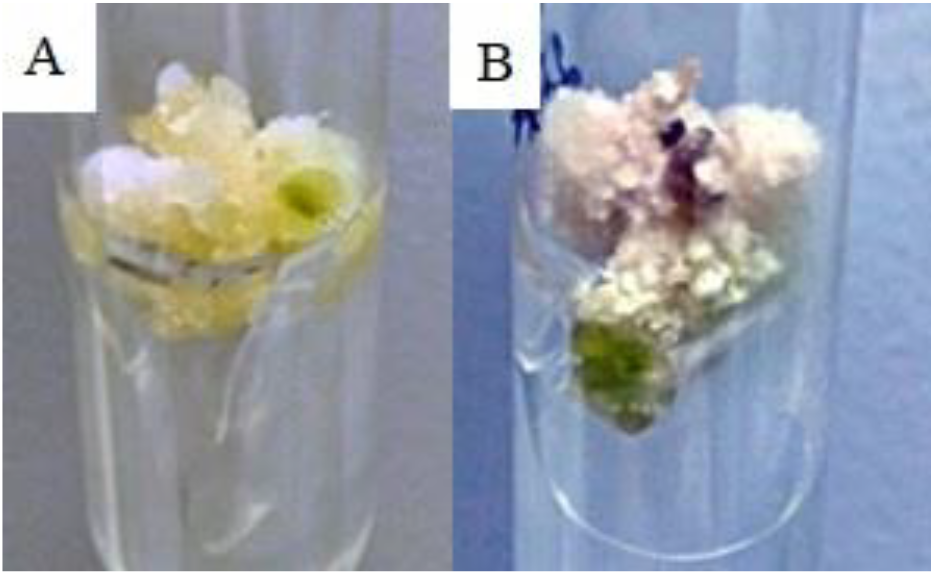
General view (A) and 25th day of growth (B) of *A. salsoloides* callus

Microscopy of meristem cells and foci of meristemoids are shown in Figure 3, respectively. The bulk of the resulting heterogeneous callus consists of meristematic type cells.

**Figure 3.**
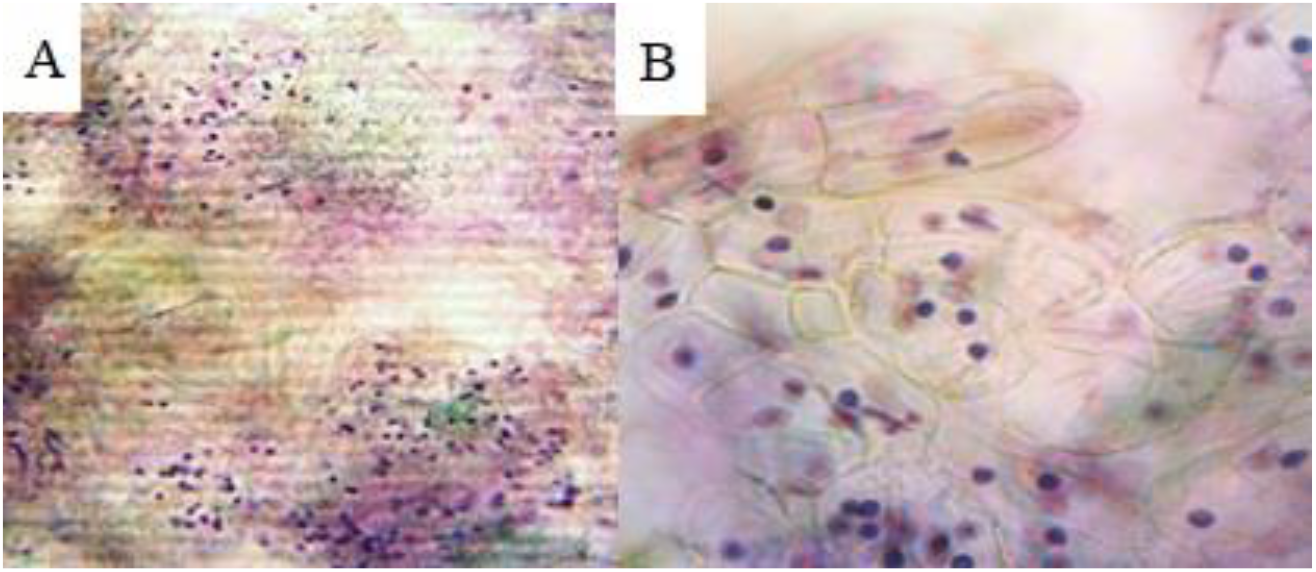
Foci of meristemoid cells (A) and meristemoid cells with stained nuclei (B) in the callus of *A. salsoloides*

After 2 weeks, in addition to callus induction, the formation of round globular structures like globular somatic embryoids (E) was noted on BM1k medium (Figure 4).

**Figure 4.**
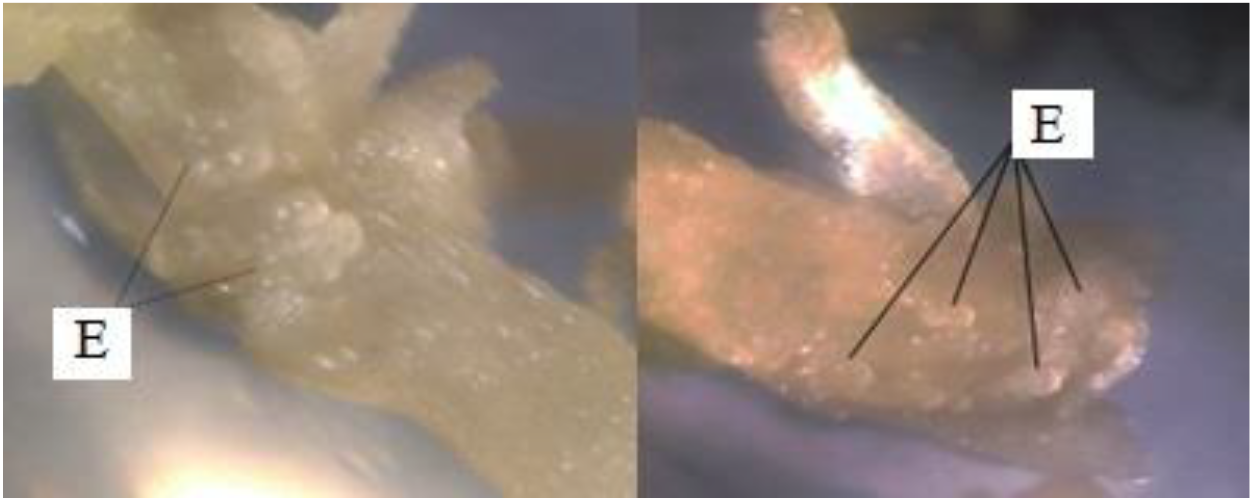
Embryoids (E) on the callus of *A. salsoloides*

After 20 days, the tubes with the callus were moved to lighting conditions, 16/8 hours-day/night. After 4 weeks, callus formation was noted on all explants (Figure 5).

**Figure 5.**
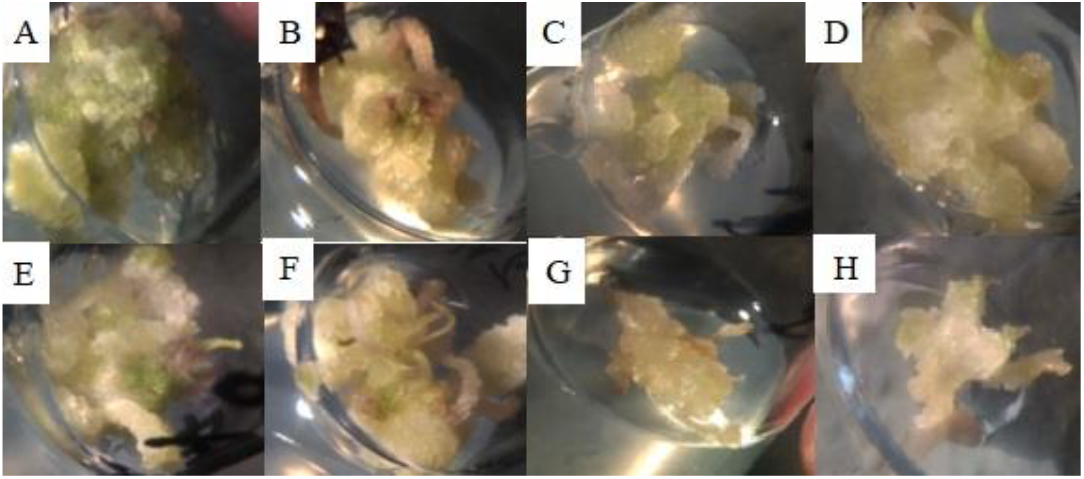
Formation of callus on BM1k (A,B,E,F) and MS (C,D,G,H) medium

In general, the callus on the BM1k medium has a denser consistency and rather has a perspective in terms of morphogenesis as a method of reproduction of red book species *in vitro*.

### Maintenance of cell suspension

Cell cultures were placed in special sterile jars with nutrient medium in a volume of 45 mL under sterile conditions of a laminar flow cabinet. Then they were tightly covered with a stretch film and placed in an orbital shaker for constant rotation (100 revolutions per minute). The suspension culture was replanted once every 4 weeks. For this purpose, 5 mL of sterile cell suspension was added to sterile jars with fresh nutrient medium (45 mL) (Figure 6).

**Figure 6.**
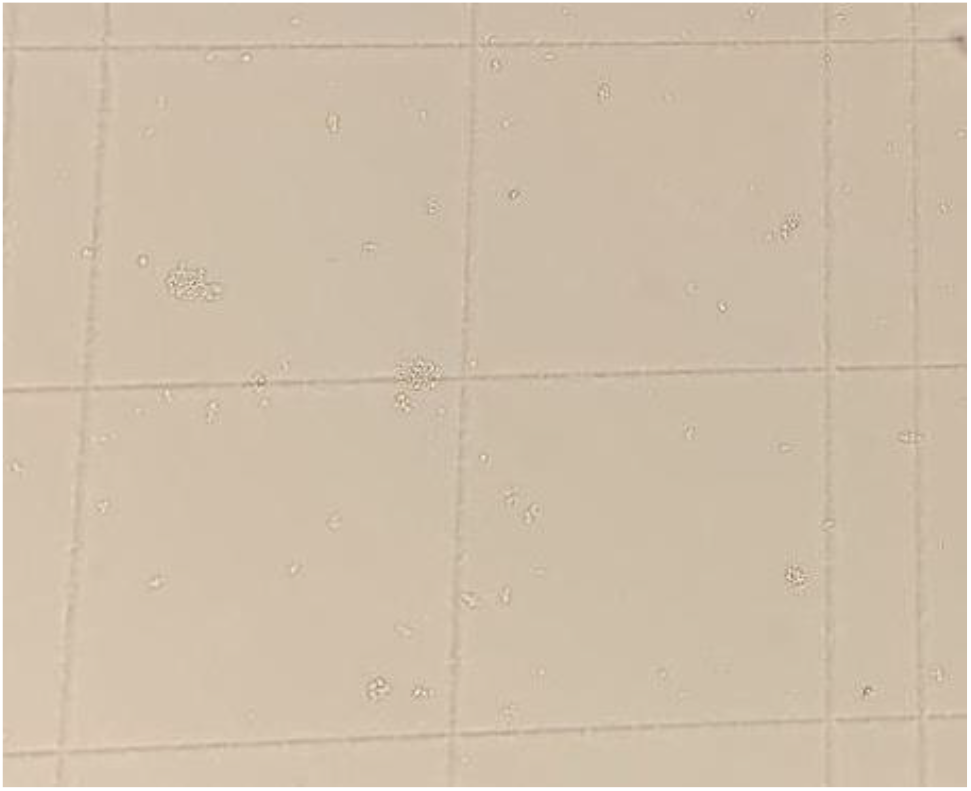
*A. salsoloides* cells in MS nutrient medium on the 14th day of cultivation.

To select the optimal nutrient medium, cells were counted using a Goryaev chamber for 2 weeks. The results obtained are shown in Table 7.

**Table 7.** Number of cells (thousand units/ml) in the suspension culture of *A. salsoloides* on 2, 4, 6, 8, 10, 14 and the 14th day of deposit.

| Nutrient media | Day |  |  |  |  |  |  |  |
| --- | --- | --- | --- | --- | --- | --- | --- | --- |
|  | 0 | 2 | 4 | 6 | 8 | 10 | 12 | 14 |
| MS | 2700±<br>571 | 3650±<br>960 | 4800±<br>1433 | 5950±<br>720 | 6500±<br>692 | 5500±<br>692 | 4650±<br>1008 | 3250±<br>692 |
| 1/2 MS | 2350±<br>1184 | 2950±<br>650 | 5100±<br>1055 | 6450±<br>571 | 7200±<br>843 | 5800±<br>1046 | 5050±<br>720 | 3900±<br>733 |

Based on the data in the table, we see that all the selected media contribute to an increase in the number of *A. salsoloides* cells for 8 days, after which we observe a decrease in the number of cells, as the nutrient medium is depleted, the amount of minerals decreases, and the number of cells is large enough, so part of the suspension culture dies.

### Genetic analysis

IPP and DMAP are involved in the synthesis of artemisin, they condense farnesyl diphosphate synthase (FPS/FPS) into farnesyl diphosphate (FPP, farnesyl pyrophosphate) (Wen and Yu, 2011). FP is then converted by amorpho-4,11-diene synthase (ADS) to amorpho-4,11-diene (AD). Next, amorpha-4,11-diene is hydroxylated to artemisinic alcohol, which is oxidized to the corresponding aldehyde, these steps are carried out by amorphadiene monooxygenase (CYP71AV1).

In addition to CYP71AV1, the oxidation of artemisinic alcohol is also carried out by alcohol dehydrogenase ADH1. At the next stage, the formed artemisinic aldehyde is reduced to dihydroartemisinic aldehyde by Δ11(13)-reductase (DBR2) and, further, oxidized by aldehyde dehydrogenase (ALDH1) to dihydroartemisinic acid. Under the action of the same enzyme, artemisinic aldehyde is oxidized to artemisinic acid. The final step is the spontaneous photoinduced non-enzymatic conversion of dihydroartemisinic acid to artemisinin and artemisinic acid to arteannoin B (Hassani et al., 2023). In its natural state, artemisin accumulates in the glandular trichomes of plants in the amount of 0.01–1.4% of dry weight (Muangphrom et al., 2016).

The study used 4 markers to assess the presence of the main artemisin synthesis genes in plants of the genus *Artemisia*. The amplification temperature (Ta) was selected experimentally. When choosing the optimal amplification temperature for each primer, we were guided by the experimental data obtained and the principle that a high annealing temperature determines a high specificity of the reaction, but as soon as it exceeds a certain critical temperature for the primer, the amount of product drops sharply.

The calculated melting point (Tm) of the primers for PCR and PCR-RV differed from the actual optimal amplification temperature (Ta). The temperature range was set from 47 °C to 57 °C and from 54°C to 62 °C, respectively. The experimentally established effective annealing temperatures for the studied primers are shown in Table 8.

**Table 8.** Optimization of temperature PCR protocols for *A. salsoloides*.

| Name of the gene | Melting temperature, (T<sub>m</sub>, °C) | Annealing temperature, (T<sub>a</sub>, °C) |
| --- | --- | --- |
| *ADS* | 65.2 | 63.2 |
| *CYP71AV1* | 65.7 | 56.2 |
| *CPR* | 64.8 | - |
| *DBR2* | 70.3 | 54.0 |

Thus, having selected the optimal ratio of PCR mixture components for each primer, the calculated melting point (Tm) and the experimentally obtained annealing temperature (Ta) did not match the calculated data, and the annealing temperature (Ta) turned out to be lower than the melting point (Tm).

However, one of the genes responsible for artemisinin synthesis, *CPR*, was not found in the species studied.

## CONCLUSION

Secondary metabolites obtained from plant cell culture *in vitro* are a promising area of work from pharmaceutical to cosmetic fields. Optimization of rare plant cultivation protocols involving secondary metabolites with potential medicinal and/or technically significant metabolites has enormous potential.

The use of standard nutrient media such as MS and its derivatives (1/2 MS) for the cultivation of sagebrush (*A. salsoloides*) has proven well with both solid nutrient media for callus formation and liquid nutrient media for the cultivation of suspension culture. However, when studying the genes responsible for producing artemisinin from wormwood plants (*A. salsoloides*) obtained from the collection “Rare and Endangered plants of the Rostov region” of the Botanical Garden of the SFedU, the *CPR* gene was not detected. Perhaps this is due to the genotype of the plants studied. In the future, natural populations will be screened and the expression of the genes responsible for the synthesis of artemisinin will be analyzed.

